# Thresholded Partial Least Squares: Fast Construction of Interpretable Whole-brain Decoders

**DOI:** 10.1101/2021.02.09.430524

**Authors:** Sangil Lee, Eric T. Bradlow, Joseph W. Kable

## Abstract

Recent neuroimaging research has shown that it is possible to decode mental states and predict future consumer behavior from brain activity data (a time-series of images). However, the unique characteristics (and high dimensionality) of neuroimaging data, coupled with a need for neuroscientifically interpretable models, has largely discouraged the use of the entire brain’s data as predictors. Instead, most neuroscientific research uses “regionalized” (partial-brain) data to reduce the computational burden and to improve interpretability (i.e., localizability of signal), at the cost of losing potential information. Here we propose a novel approach that can build whole-brain neural decoders (using the entire data set and capitalizing on the full correlational structure) that are both interpretable and computationally efficient. We exploit analytical properties of the partial least squares algorithm to build a regularized regression model with variable selection that boasts (in contrast to most statistical methods) a unique ‘fit-once-tune-later’ approach where users need to fit the model only once and can choose the best tuning parameters post-hoc. We demonstrate its efficacy in a large neuroimaging dataset against off-the-shelf prediction methods and show that our new method scales exceptionally with increasing data size, yields more interpretable results, and uses less computational memory, while retaining high predictive power.

---

Functional magnetic resonance imaging (fMRI) has allowed researchers to empirically examine the relationship between brain activity and various behaviors and cognitive processes. Brain activities are measured in small grids, in 3D units known as voxels (i.e., volumetric pixels), and a typical fMRI image of the brain can have anywhere between 30,000 and 1.8 million voxels (4mm and 1mm resolution respectively). The number of images per person is typically much smaller, in the hundreds, and the number of people who participate in a study is smaller still, typically less than a hundred. This is because data needs to be collected in person, while conducting an experimental task inside an MRI machine that is operated by a trained technician. Therefore, the regime of predicting cognitive states and behavior using voxels in fMRI is in the domain of classic large *P*, small *N* problems (*P* ≫ *N*), where the number of predictors is much larger than the number of observations.

To address the large *P*, small *N* problem, most fMRI studies have taken one of two main approaches. The first approach, which the bulk of fMRI studies have taken, is to avoid the problem altogether, by focusing instead on predicting neural activity from measured behavioral or cognitive states, usually in a “massively univariate” manner that considers each single voxel independently in parallel (Friston et al., 1994; S. M. Smith et al., 2004). In the second approach, those studies that have tackled predicting behavioral and cognitive states from neural activity have usually focused on partial-brain prediction from small cutouts of the brain (lowering P). For example, a common method is “searchlight analysis”, in which multivariate predictors are built taking a handful of local voxels at a time, iterating through the brain to identify which regions can predict the given behavior or cognitive states above chance (Etzel, Zacks, & Braver, 2013; Kriegeskorte, Goebel, & Bandettini, 2006).

These approaches have served neuroscientific research well, as often the goal of an fMRI study is to test *a priori* hypotheses about the function of specific brain regions. That is, for the neuroscientists, it’s less important that prediction of behavior is possible than to know which brain area makes it so. However, there are several reasons why one may still want to predict behavioral or cognitive states using *all* the voxels in the brain (for review, see Kragel, Koban, Barrett, & Wager, 2018; Woo, Chang, Lindquist, & Wager, 2017). First, from a neuroscience perspective, a whole-brain predictor can give insight into the relationship between the signals in different brain regions, much like how multiple regression can give different answers than pairwise correlations. Second, from the perspective of psychologists or others seeking to apply neuroscience methods to problems in other domains, prediction *per se* is often the primary goal. For example, one can use brain predictors as biomarkers for certain disabilities or impairments. In this case, a whole-brain predictor can provide higher sensitivity than regional methods by combining signals across the entire brain, and can also provide higher specificity by identifying a unique predictor for each mental state, since the response pattern in any given region of the brain may occur with many different mental states.

Given these benefits, there have been several recent efforts to construct generalized whole-brain decoders. For example, Wager et al. (2013) has constructed a whole-brain signature of pain that predicts the degree of pain an individual feels. Smith et al. (2014) and Lee, Lerman, & Kable (2019) have constructed whole-brain predictors of valuation. Chang et al. (2015) constructed a whole-brain neural signature of picture-induced negative affect; Kragel & LaBar (2014) and Kassam, Markey, Cherkassky, Loewenstein, & Just (2013) constructed neural predictors of distinct emotional states.

Unfortunately, these efforts building whole-brain decoders with off-the shelf statistical methods for high-dimensional data have run into two big difficulties. A first difficulty is interpretability: a good whole-brain predictor should distinguish *regions* that are predictive versus those that are not. As a counterexample, predictors built from LASSO, a common penalized regression approach, result in predictive maps with scattered sparkles of coefficients across the brain rather than any interpretable clusters or regions. This is because while LASSO selects the most useful variables (voxels) for prediction, it does so without regard for spatial contiguity of the variables (Grosenick, Klingenberg, Katovich, Knutson, & Taylor, 2013). On the other hand, other methods often result in no variable selection such that the entire brain has non-zero coefficients. These include ridge regression (Grosenick et al., 2013), support vector machines (Whitehead & Armony, 2019), partial least squares (McIntosh, Bookstein, Haxby, & Grady, 1996), or PCR-LASSO that uses PCA for data reduction and then LASSO regression to select the most useful components (Wager et al., 2013). These approaches typically use the entire brain’s coefficients for prediction, but later threshold the coefficients, at an arbitrary level or by bootstrap, to improve interpretability in inference (McIntosh et al., 1996; Wager et al., 2013). Hence, the voxel selection is not driven based on cross-validation. One method that does yield interpretably clustered coefficients is GraphNet, which combines the elastic net penalty with voxels’ spatial contiguity information (Grosenick et al., 2013).

However, there is also a second difficulty, which is computational efficiency. This difficulty is particularly acute with regards to scaling for use in larger datasets, the collaborative collection of which is an increasing focus of fMRI research (Allen, Sudlow, Peakman, & Collins, 2014; Bjork, Straub, Provost, & Neale, 2017; Satterthwaite et al., 2014; Van Essen et al., 2012). Since neuroimaging data has a substantial number of predictors, purely likelihood-based approaches often face the problem of calculating gradients for a large number of variables. Adding to the burden, modern models need to be fitted multiple if not hundreds of times to find the best tuning parameters (e.g., the tuning parameter *α* in LASSO that controls variable selectivity). These problems are computationally challenging even in average sized fMRI datasets, which is why previous whole-brain predictors used down-sampled images (i.e. coarser) with fewer voxels for prediction (e.g., Grosenick et al., 2013; Wager et al., 2013). As an alternative to purely likelihood-based methods, data-reduction approaches such as PCA can help in immensely reducing the number of variables and thereby reducing the model fitting time. However, in larger datasets, PCA itself can become a bottleneck for computation time and memory usage as it requires computation of the variance-covariance matrix of predictors. These computational costs prevent the widespread use of whole-brain prediction methods in neuroimaging, especially for those without access to high performance computing clusters.

Here we propose a novel method, thresholded partial least squares (T-PLS, pronounced Tea, Please), that provides interpretable whole-brain predictors that are computationally efficient enough to run on personal computers for most datasets. T-PLS exploits analytical properties of a modified partial least squares algorithm to offer a unique ‘fit-once-tune-later’ approach where the user fits the model only once and then evaluates the best tuning parameter as many times as needed without re-fitting the model. This is in stark contrast to most, if not all, modern methods that require re-fitting the model for every tuning parameter. Here, we describe the algorithm and showcase its performance against other methods in a large neuroimaging dataset. Furthermore, we provide the T-PLS package online for MATLAB at github (https://github.com/sangillee/TPLSm), for R at CRAN (https://CRAN.R-project.org/package=TPLSr), and for python at github (https://github.com/sangillee/TPLSp) as (we hope) a practical tool for others.

## Methods

### Overview

T-PLS, like many modern regression methods, requires two estimation steps when training a model: fitting and parameter tuning. In the fitting step, the end goal is to calculate the coefficient and the z-statistic of each original variable (e.g., voxel) – similar to the coefficients and t-statistics one would get for each variable in multiple regression (**Fig. 1**). In detail, the fitting step first extracts the PLS components that maximally explain the covariance between X (e.g., brain voxels) and Y (e.g., behavioral or cognitive state) and then regresses Y against the components1. The resulting coefficients and the t-statistics of the components are then back-projected into the original variable space to calculate a coefficient and a z-statistic for each voxel. In the tuning step, two parameters are chosen: the number of PLS components used and the voxel z-statistic threshold that controls the number of voxels retained. For example, using cross-validation, a T-PLS model that uses the first 6 PLS components and retains 50% of the original voxels may provide the highest out-of-sample cross-validation performance. Intuitively, the number of PLS components controls the degree of data reduction while the voxel thresholding level controls variable selection, and, as desired here, improves interpretability.

**Figure 1.**
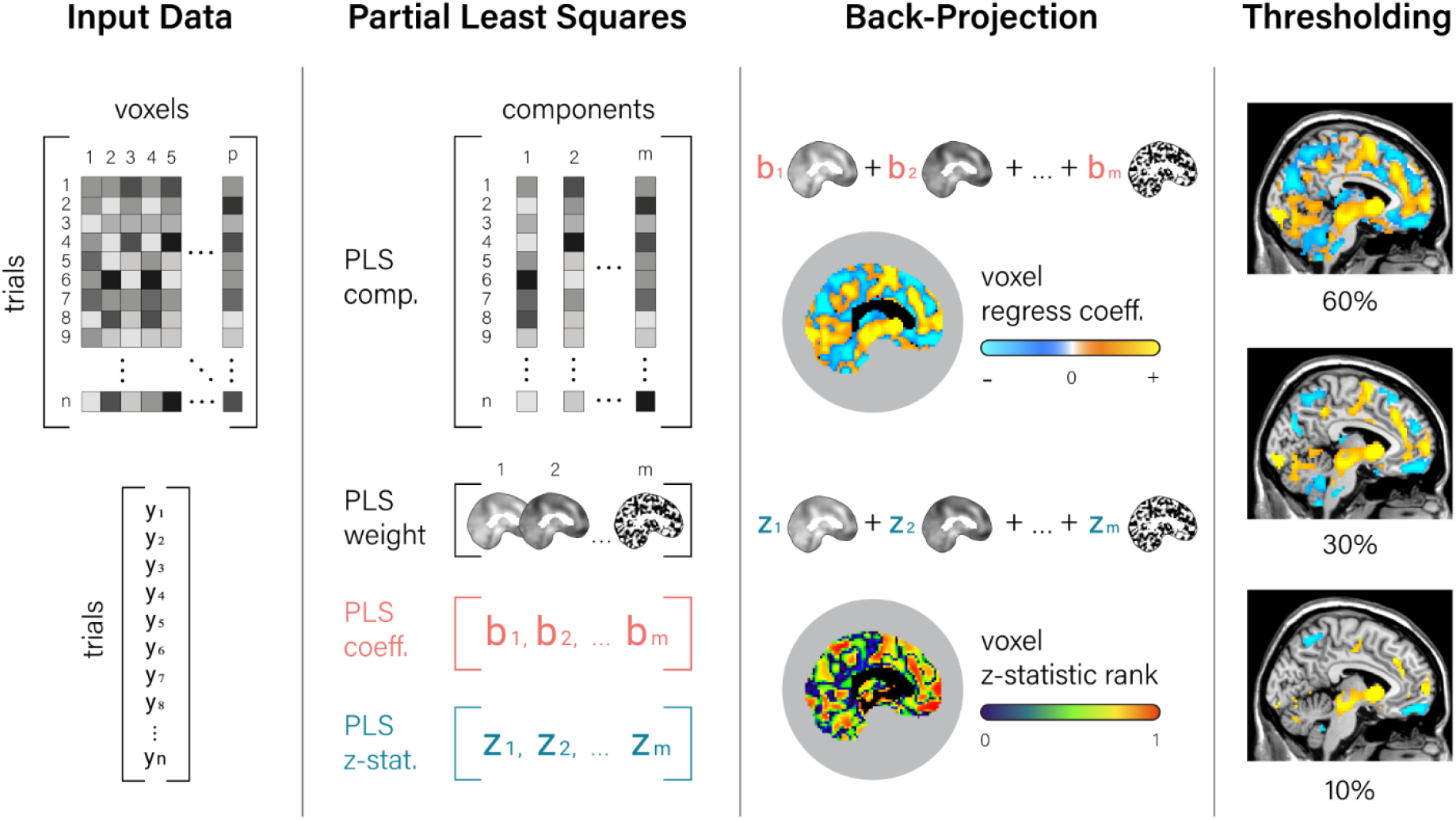
Schematic of T-PLS fitting procedures. T-PLS model fitting first requires extracting the partial least squares components from the predictor matrix and obtaining the components’ back projection maps (PLS weight), their regression coefficients, and their z-statistic. Next, the regression coefficients and the z-statistics are back-projected into the voxel space using the weight maps, thereby yielding a whole-brain coefficient map and a whole-brain z-statistic map, which is then ranked in absolute size to determine voxel importance. Finally, the coefficient map is thresholded based on the z-statistic rank map to select voxels that are the most important.

The key computational benefit of T-PLS comes from the ‘fit-once-tune-later’ feature. The user can choose among infinitely many tuning parameter combinations without having to re-fit the model, as all the information required is already calculated in a one-time fitting. This is because once a T-PLS model with *m* components has been fit, all the models with fewer components are also available (i.e., a 1-component, 2-component, …, *m*-component model). Since PLS components are all orthogonal to each other, their regression coefficients do not change based on the number of components kept, thereby allowing the user to choose the necessary components without re-fitting.

Using PLS for data reduction (as in T-PLS) provides two additional key benefits over PCA (as in PCR-LASSO). First, PLS computation only requires vector multiplications, which are fast and memory efficient, while PCA requires matrix singular value decompositions, which grow quadratically with data. Second, PLS components are ordered in terms of the explained covariance of X on Y, while PCA components are ordered only in terms of variance explained in X. A standing criticism of using PCA for data reduction is that while PCA components are great for describing X, they are not necessarily relevant in predicting Y (Lever, Krzywinski, & Altman, 2017).

### T-PLS Algorithm - fitting

The fitting algorithm for T-PLS is a combination of three parts: a modified SIMPLS algorithm for PLS (de Jong, 1993), back-projection, and calculation of z-statistics (**Fig. 2).** The one modification that we make to the SIMPLS algorithm is simply in normalizing the PLS components to have weighted unit variance (step 4 in **Fig. 2**). This facilitates computation of z-statistics later in the algorithm (step 7 in **Fig. 2**). Below we detail the back-projection and z-statistic calculation steps of T-PLS as the remaining steps are typical procedures of a SIMPLS algorithm.

**Figure 2.**
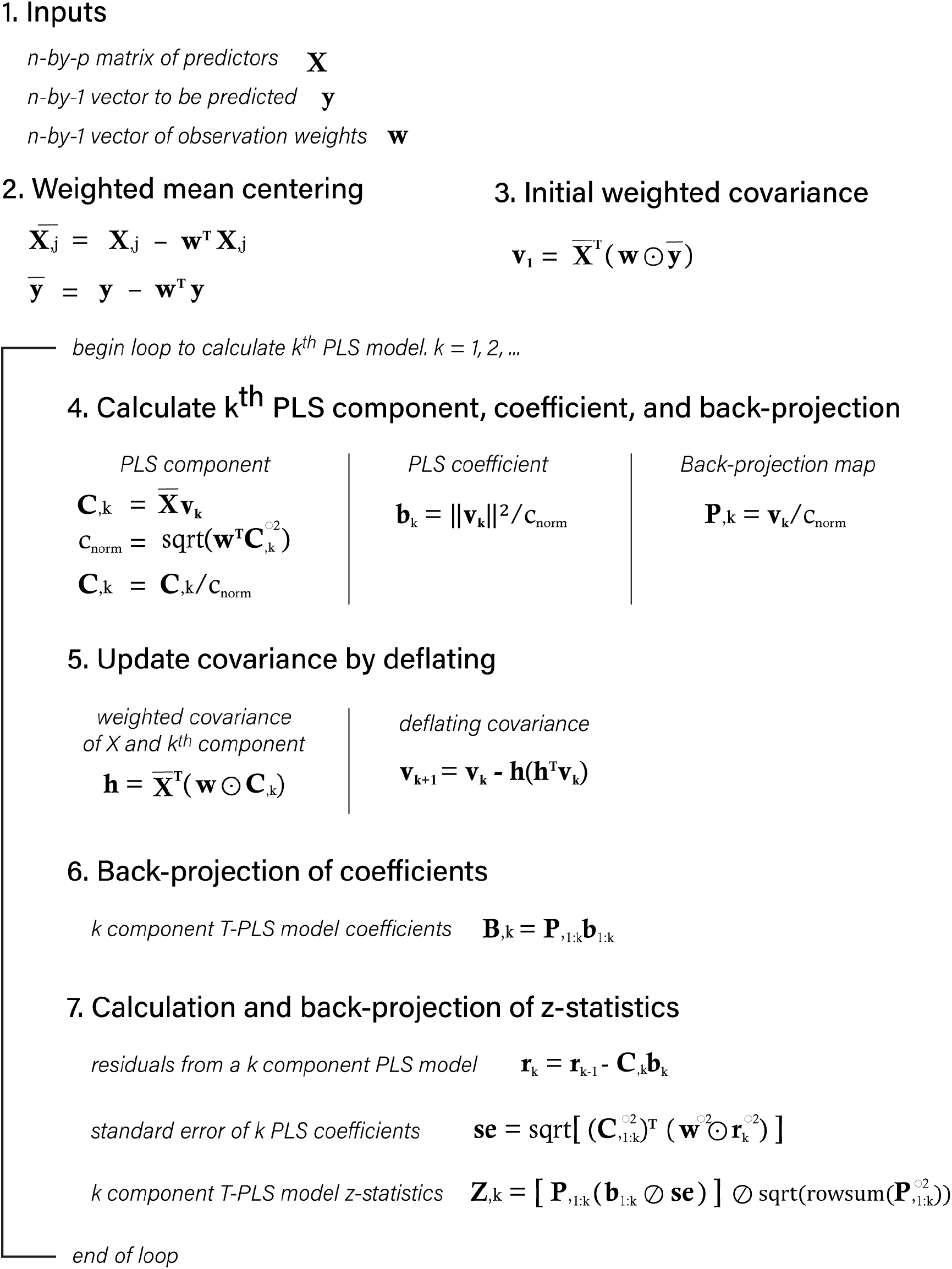
Summary of T-PLS fitting algorithm. Matrices are denoted with bold capital letters, vectors with bold lowercase letters, and scalars with non-bolded lowercase letters. ⊙ denotes Hadamard product (elementwise multiplication), ⊘ denotes elementwise division, and ∘ 2 in the exponent denotes elementwise squaring.

*Back-projection* (step 6 in **Fig. 2**). After PLS components have been calculated (up to *k* ^th^ component), we now have a *k*-component PLS regression model. To improve the interpretability of this PLS model, we can convert the PLS regression coefficients into coefficients for the original voxels. Since PLS components are created via weighted sums of original voxels (i.e., component = weight * voxels), one can simply multiply the PLS coefficient to the weights to create back-projected coefficients (i.e., coefficient * component = coefficient * weight * voxels = back-projected coefficient * voxels). This expresses the PLS regression in terms of each voxel’s coefficients, which can make the predictor easier to interpret by identifying which regions are positively or negatively predictive of behavior or mental states. Back-projection is also used in the PCR-LASSO method applied by Wager et al. (2013), but with PCA components rather than PLS.

*Z-statistic calculation* (step 7 in **Fig. 2**). We then calculate the z-statistic of each voxel as a measure of variable importance. We start by calculating the heteroscedasticity-consistent standard errors (also known as sandwich estimators; White, 1980):

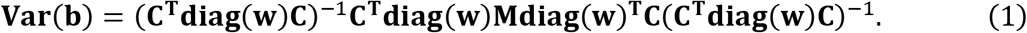

Where **b** denotes the coefficient estimates, **w** denotes the observation (trial) weights, **M** denotes the variance-covariance matrix for the observations, and **C** denotes the PLS components in a column-wise matrix. Here is where our modification to the SIMPLS algorithm becomes useful. Since the PLS components (matrix **C)** are all orthonormal (in weighted space), **C** _**T**_**diag(w)C** becomes an identity matrix, which cancels out the ‘breads’ of the sandwich and leaves us with **Var(b)** = **C**_**T**_**diag(w)Mdiag(w)**_**T**_**C**. Since we only need the diagonals of the variance-covariance matrix, we can express the standard error estimates concisely as the following:

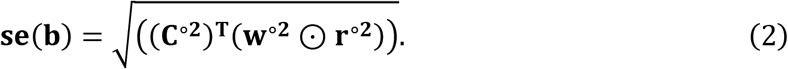

where **r**^**∘2**^ denotes the squared residual vector. The t-statistics (which are close to z-statistics with sufficient observations) can be then calculated by simple element-wise division of *b* by **se(b)**. Let this vector be denoted **z**. Then, we back-project the z statistic like the coefficients, and then normalize them so that they all have unit variance:

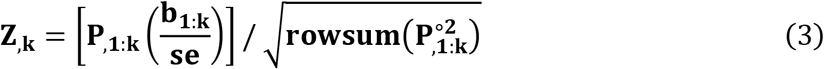

where **Z**_**,k**_ is the z-statistics of each voxel calculated from a *k* component T-PLS model, and rowsum denotes the vector that is the row sum of a matrix. This summarizes the fitting procedure of T-PLS.

### T-PLS Algorithm - tuning

After the model fitting is complete, one can retroactively choose the two tuning parameters, the number of PLS components and the voxel thresholding level, based on cross-validation performance. **Fig.3** shows an example cross-validation performance surface as a function of the number of PLS components and the voxel thresholding level. Because the model does not have to be re-fit, researchers can examine the predictor at various thresholding levels and assess the tradeoff between predictive performance and model sparsity.

**Figure 3.**
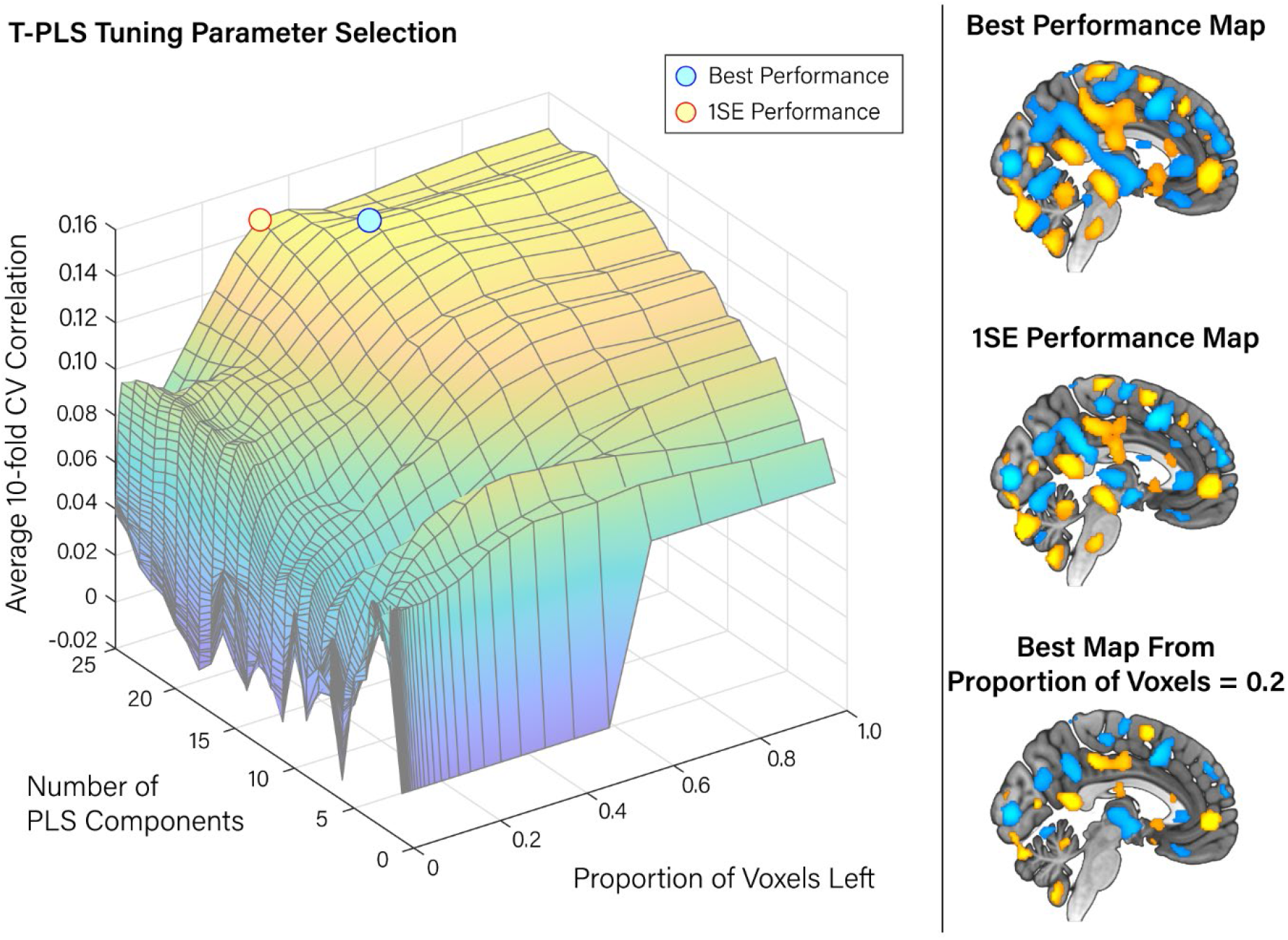
Example T-PLS model tuning. Left panel shows an example cross-validation performance surface as a function of the two tuning parameters of T-PLS: number of PLS components (1~25) and proportion of voxels left (0~1). The highest CV performance point is marked with a blue dot with the corresponding whole-brain predictor shown on the right top panel. Additionally, a model with fewer voxels but within 1 standard error of the best model’s performance is indicated with a yellow dot with the corresponding map shown on the right middle panel. The right bottom panel is shown to illustrate how the number of remaining voxels with coefficients reduce as the proportion of voxels left are reduced.

After the tuning is complete, there is one more step that may be useful in some scenarios: post-fitting of bias (intercept). Since some variables are removed during the thresholding stage, the intercept should be re-fitted after thresholding. Let’s say that we chose to evaluate a model with *j* components, thresholded at 70% (removing 70% of variables). Then, the coefficients are **B.**_**,j**_ multiplied by index vector **d**where **d**_**i**_ = 1 if the voxel’s rank is in the top 30% and 0 otherwise. Then the new intercept is simply the difference between the weighted means of **X** and **y**:

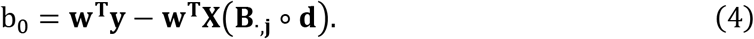

### Neuroimaging Dataset

We use a large neuroimaging dataset from Kable et al. (2017) to empirically compare T-PLS against two other whole-brain methods (PCR-LASSO and LASSO) as well as region-based (partial-brain) predictions. We chose this dataset as it was the most readily available large-scale dataset with whole-brain coverage that can be used to decode behavior from brain activity levels. Participants completed two experimental decision-making tasks, intertemporal choice and risky choice, which are both very common in the domain of social science (psychology, economics research, e.g., Green & Myerson, 2004; Kahneman & Tversky, 1979; Samuelson, 1937) and its interaction with neuroscience (e.g., Jung, Lee, Lerman, & Kable, 2018; Kable & Glimcher, 2007). In intertemporal choice, participants made choices between a smaller immediate monetary amount of $20 and a larger but delayed monetary amount (e.g., $40 in 30 days). This choice paradigm allows one to assess the “time value of money”, hence its name “intertemporal choice”. In risky choice, subjects made choices between a smaller certain monetary amount of $20 and a larger but probabilistic monetary amount (e.g., $40 with 60% probability of winning). In both tasks the larger amount varied from trial to trial, as well as the associated delay or the risk, while the smaller monetary option was always fixed at $20. Only the larger monetary option was on the screen while the smaller $20 was not; participants made accept/reject choices based on whether they would prefer the larger monetary option on the screen or the smaller monetary option. Because the value of one of the options was always constant, Kable et al. (2017) were able to find signals in the brain that correlated with the subjective value of the varying option that was shown on the screen. Based on this result, we seek here to create a whole-brain predictor of choice that can use these signals to predict whether the participant will accept the option on the screen or reject it. Details about removed participants, fMRI image acquisition protocols and preprocessing details are provided in the **supplemental materials**. In total, the dataset gave us a total of 61,038 trials (observations) and 184,319 voxels (variables) across 531 task sessions (264 intertemporal choice sessions, 267 risky choice sessions), which we treat as 531 participants in this paper, as our goal is not in making substantive, or comparative, conclusions about the tasks.

### Computation comparisons

We assess the scalability of each whole-brain prediction method – LASSO, PCR-LASSO, and T-PLS – by comparing their model fitting time and RAM usage at varying training dataset sizes (8, 16, 32, 64, 128, 256, and 512 participants). In each dataset size (e.g., 8 subjects), half of the data is drawn randomly from the risky choice dataset (i.e., 4 subjects) and the other is drawn randomly from intertemporal choice dataset. Each model is fitted using 10-fold cross-validation (CV). The training data is divided into 10 equal sized blocks and the model is fitted on 9 of the blocks and tested on the left-out block. This is repeated 10 times to assess cross-validation performance. Then, the tuning parameter that yields the highest CV performance is chosen and used to train the final predictor using all training data.

For LASSO, we use GLMNET for MATLAB (Friedman, Hastie, & Tibshirani, 2010; Qian, Hastie, Friedman, Tibshirani, & Simon, 2013), which is arguably the fastest non-GPU package for fitting LASSO thanks to its use of regularized path and FORTRAN coding^2^. We use the default tuning parameter search, which uses 100 lambda values. For PCR-LASSO, we stick with the original approach in the Wager et al. (2013) paper by extracting 200 components from all data, using 10-fold LASSO logistic regression to find the useful components, and subsequently running an unpenalized logistic regression using only the selected components. For T-PLS, in each of the 10 folds, we extract 25 PLS components and built the T-PLS model. Then, during cross-validation we choose the best-performing number of PLS components and threshold level. Each whole-brain method is fitted 400 times at each dataset size, each time randomly selecting the training data. All computations were performed on a large-scale computation cluster at the University of Pennsylvania (https://www.med.upenn.edu/cbica/cubic).

### Predictive power comparisons

We compare the out-of-sample predictive performances of the predictors built above. After the prediction model is fitted using 10-fold cross validation in the training data (e.g., 32 subjects), the remaining data (e.g., 531-32=499 subjects) is used as an out-of-sample testing dataset. Per-subject correlation and area under the ROC curve (AUC) are averaged across the out-of-sample participants to get an estimate of out-of-sample prediction performance. We also add two commonly used region-based prediction methods to the comparison of predictive power: region-average, and region-multivariate. Region-average is simply taking the average of all voxel activities within a designated region to make predictions; concordantly, region-average does not require fitting a model. Region-multivariate, on the other hand, uses the voxels in the region to build a predictor. While several methods can be used, here we use LASSO to make comparisons with our whole-brain methods easier. We use regions identified from a meta-analysis by Bartra, McGuire, & Kable (2013), which examined around 150 neuroimaging studies and identified two regions that consistently showed correlated activity with valuation: ventral striatum and ventromedial prefrontal cortex (regions from figure 9 of Bartra, McGuire, & Kable 2013).

### Interpretability comparisons

We also compare the interpretability of the three whole-brain methods (LASSO, PCR-LASSO, T-PLS). Using the same fitting procedures as before (10-fold cross-validation), T-PLS and PCR-LASSO are fit using the entirety of the data (531 participants). LASSO, however, is only fit with a subsampled 64 participant dataset, as it is computationally too slow to fit using the entire dataset. We visually compare the resulting whole-brain predictors and the associated areas of the brain to assess the scientific face validity of the identified brain regions.

Additionally, we use simulated data to provide further insight into differences in interpretability. We simulate a brain activity signal of a 17×17 voxel grid (total of 289 voxels), of which only a 5×5 grid in the center (25 voxels) carries signal that is predictive of Y, while all other voxels are completely orthogonal to Y (i.e., noise). We achieve this by first randomly generating 290 variables (289 voxels + 1 Y) each with 300 observations from a standard normal distribution. Then, we apply symmetric orthogonalization such that all 290 columns are orthogonal to each other. The first column of the new matrix is chosen as the predicted variable **Y**, while the other 289 variables became simulations of fMRI noise. To create 25 voxels of predictive voxel signal, we mix **Y** with 25 of the simulated fMRI noise variables to create 25 signals that are all exactly correlated with Y at r = 0.1. Each column is then z-scored to have unit variance. Finally, we place the 5×5 signal grid in the center of a 17×17 grid and apply 2D Gaussian smoothing (sd = 1 voxel) to simulate the inherent smoothness of fMRI signals. In sum, the resulting dataset is 300 observations of 17×17 voxel grid predictors with only the center 5×5 grid being predictive of Y. This simulated dataset is fit by OLS, LASSO, PCR-LASSO (10 components), and T-PLS models (10 components) to compare the resulting pattern of coefficients.

## Results

### Computation Time

T-PLS shows exceptionally fast model-fitting time that scales very easily to large datasets (**Fig. 4A**). In the largest training dataset size of 512 people (256 sessions of ITC and 256 sessions of risky choice), T-PLS took 2 hours and 10 minutes on average to finish 10-fold cross validation training. PCR-LASSO was 28 times slower than T-PLS, taking 2.3 days. LASSO was already taking close to 2 weeks for 256 participants and was too expensive to compute for larger dataset sizes. Importantly, when the fitting time per-participant was assessed, T-PLS was the only algorithm that maintained a constant model fitting speed (~15 seconds), whereas PCR-LASSO and LASSO show increasing fitting time per-subject as dataset size increases (**Fig. 4B**). For PCR-LASSO, this increase in fitting time per-subject is likely because PCA requires inversion operations on the variance covariance matrix of X, which quadratically increases in size until the number of observations matches the number of variables. For LASSO, this increase in fitting time per-subject is also likely due to calculation of gradients based on matrix operations. T-PLS, in contrast, only requires vector calculations, for which only the number of variables is the dominating factor. The speed of T-PLS will prove useful for large-scale neuroimaging studies as well as studies that may train many different predictors for different behaviors or mental constructs. In addition, since T-PLS only has to be fit once, this will provide an even greater benefit in computational time.

**Figure 4.**
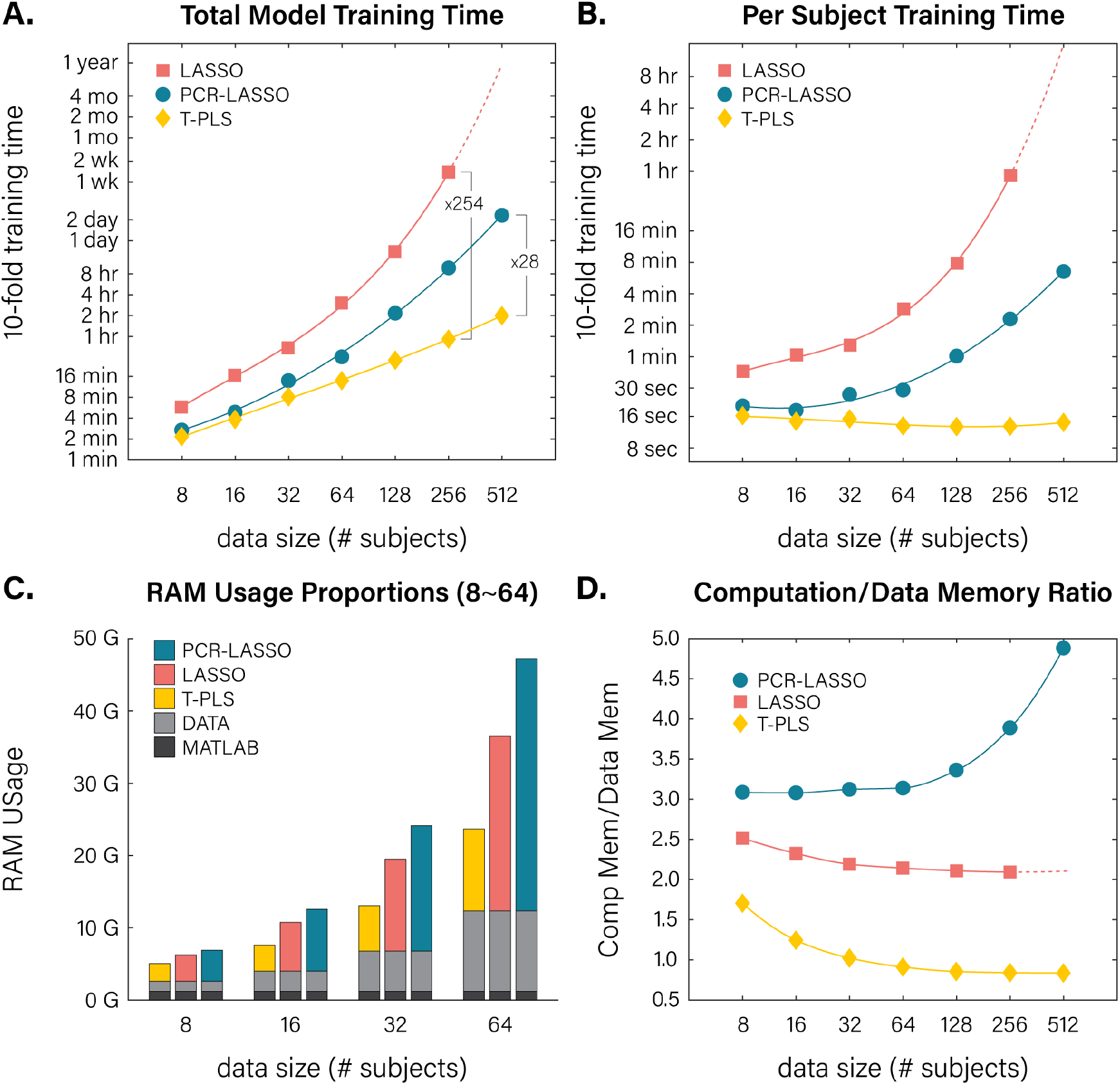
Computation resource comparison. Panels A and B show 10-fold model training times for each algorithm at various dataset sizes (total time and total time dived by number of subjects, respectively). The lines show best fitting cubic polynomial spline fit. Panels C and D show RAM usage of each algorithm at various dataset sizes. Panel C shows the memory decomposition of each algorithm (memory for turning on MATLAB, for loading data, and for computation). Panel D shows the ratio between RAM usage for loading data and RAM usage for computation (colored vs. light grey bar in panel C).

### Memory Usage

T-PLS also uses a very minimal amount of memory compared to other algorithms. (**Fig. 4C**). We broke down memory usage into three parts: default RAM for loading the statistical program, RAM for loading the data, and RAM for computing the model from the data. The first two parts are the same across all algorithms as they all need to load the program and the data. The differences across algorithms come from the differences in RAM usage for model computation. T-PLS used the least amount of computation memory, followed by LASSO and PCR-LASSO.

We again found that T-PLS was the most scalable out of all three algorithms (**Fig. 4D**). T-PLS’s computation RAM usage converges to about the same amount as RAM needed for loading the data (1.75 times -> 0.95 times as dataset size increases). This is likely because T-PLS requires a mean-centered copy of the data matrix. Should researchers want, they can mean-center the data beforehand and use even less memory for T-PLS (this feature is available in the provided statistical packages). In contrast, LASSO’s RAM usage for computation ranged from 2.6 times to 2.2 times the data RAM size, while PCR-LASSO’s RAM usage for computation increased rapidly as dataset size increased (3 times -> 4.9 times).

### Prediction Performance

T-PLS showed the highest predictive performance across all tested dataset sizes (**Fig. 5**), with the caveat that LASSO was too expensive to compute at the largest dataset size, since it was expected to take one year on average to fit. All three whole-brain predictors provided predictive performances that increased from the smallest dataset size to the largest dataset size. T-PLS showed the highest predictive performances across all dataset sizes. LASSO’s performance was the worst out of the three whole-brain methods in small dataset sizes, but increased rapidly to be better than PCR-LASSO. There were also significant differences between whole-brain methods and region-based methods. Predictors based on only the voxels within a pre-defined region provided the worst performances across the board, with the region-multivariate method outperforming the region-average method. This demonstrates how a whole brain predictor can harness more signals from across the brain to provide greater predictive power.

**Figure 5.**
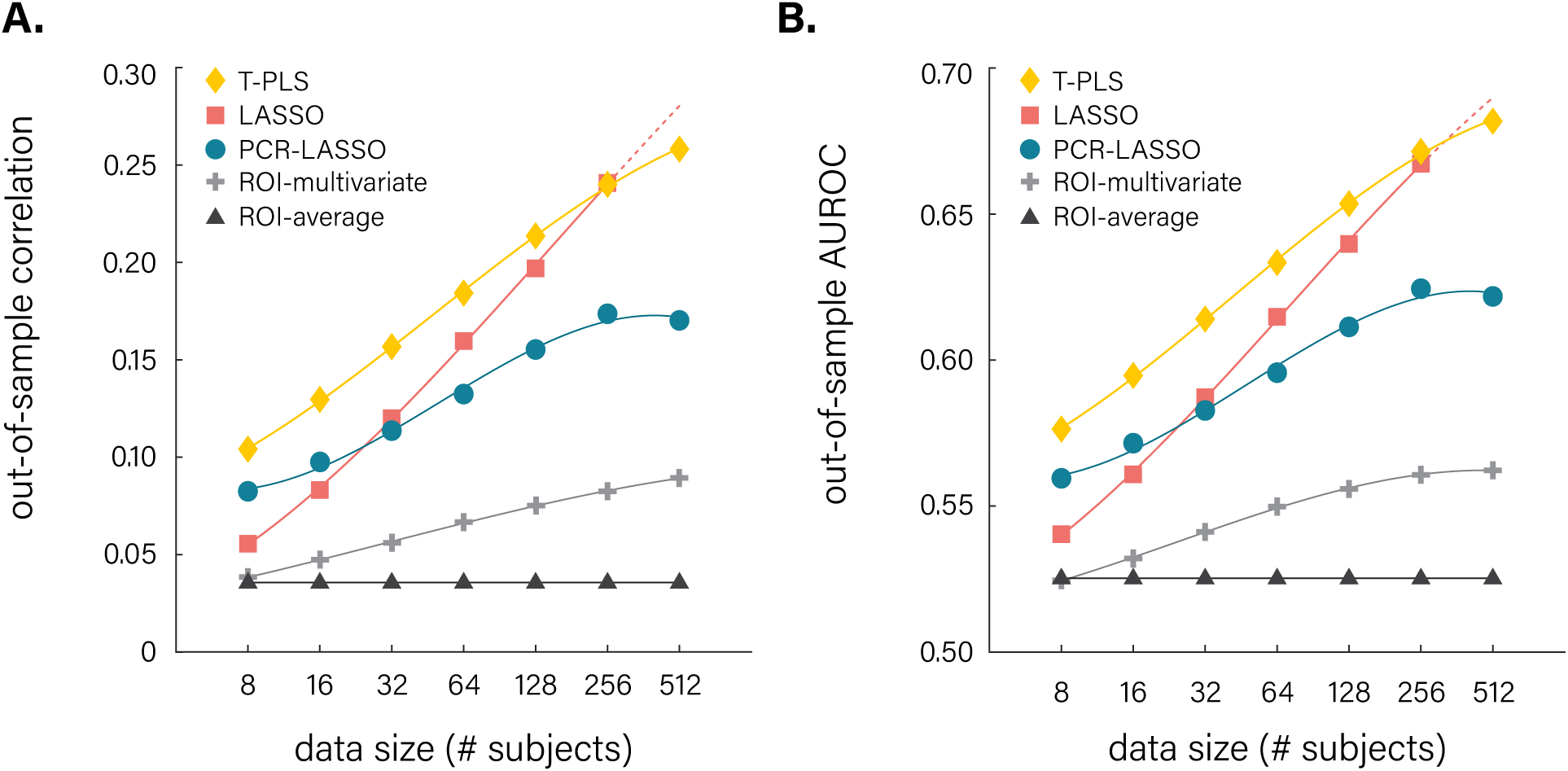
Out-of-sample prediction accuracy of various algorithms. Shown above are out-of-sample prediction performances of five algorithms measured via Pearson correlation (A) and AUROC (B) for predicting value-based accept/reject choices. The lines show best fitting cubic polynomial spline fit.

### Predictor Interpretability

T-PLS provides predictor maps that are easily interpretable and that differ in important ways from those of other approaches (**Fig. 6**). T-PLS results in whole-brain predictors with regionally clustered coefficients and voxel selection (**Fig. 6A**). This allows researchers to identify key brain regions in the predictor. PCR-LASSO also leads to regionally clustered coefficients, but includes no voxel selection (**Fig. 6B**). In contrast to the other two methods, LASSO predictors select single voxels that are most important for prediction; however, single voxels can be very difficult to interpret as it is not always easy to pinpoint the region from which the voxels originate (**Fig. 6C**).

**Figure 6.**
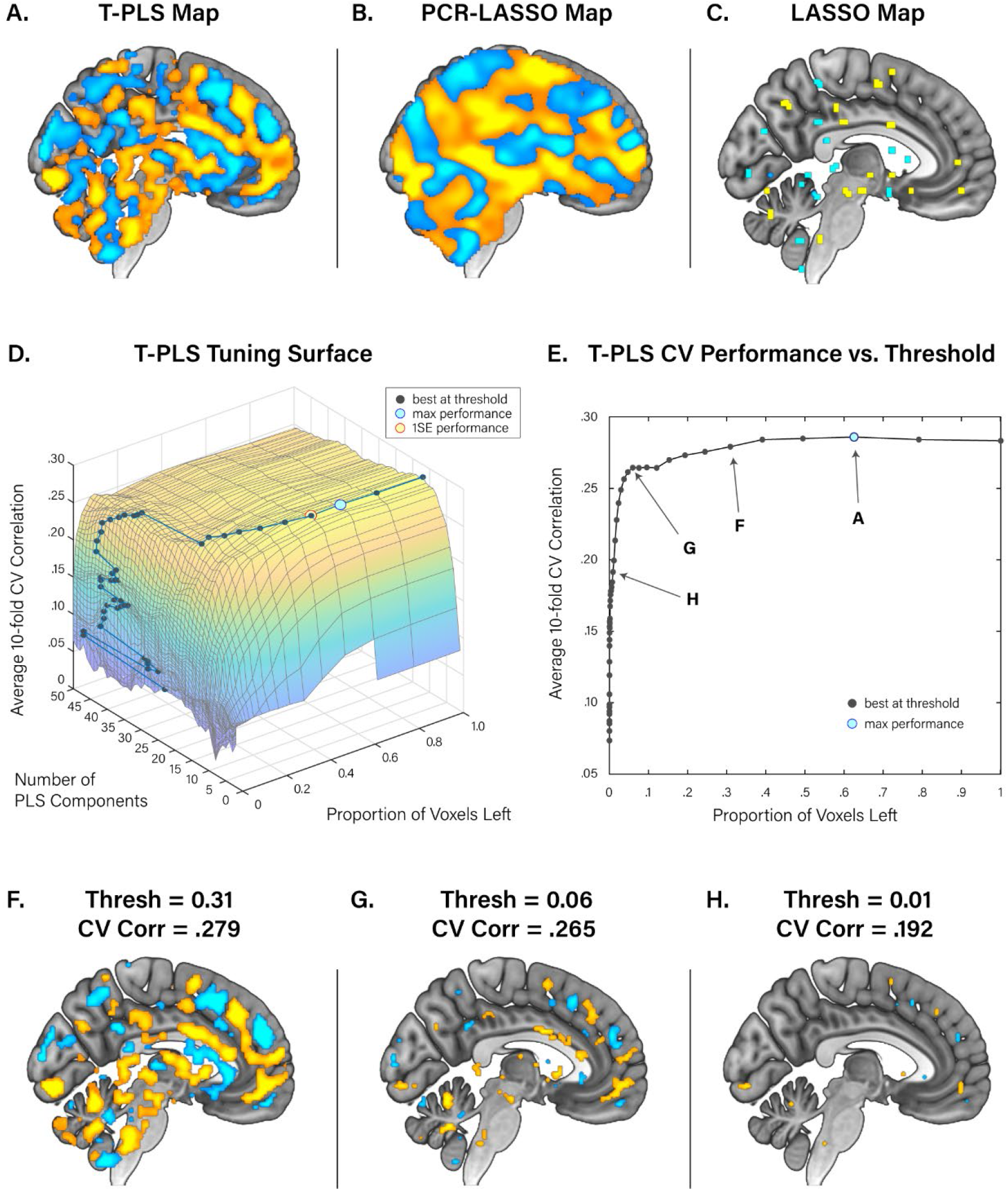
Final predictors of value-based choice. Panels A, B, and C show the whole-brain predictor of value-based choice constructed via T-PLS, PCR-LASSO, and LASSO, respectively. Panel D shows the 10-fold cross validation tuning curve for fitting the T-PLS predictor. Panel E shows the 10-fold cross validation performance of the T-PLS model at various thresholding levels. Panel F, G, and H show the corresponding thresholded predictors from panel E.

Apart from the interpretability of the finished predictor, T-PLS can also provide useful information on the relative importance of different brain regions by showing the tradeoff between additional thresholding and cross validation performance (**Fig. 6D & E**). Users can experiment with different thresholds to see how much predictive performance is sacrificed when fewer regions are recruited into the predictor. **Fig. 6F~H** shows various predictors at different thresholds where stringent thresholds (e.g., **Fig 6H**) highlight the more important brain regions for prediction. This analysis is possible due to T-PLS’s fit-once-tune-later approach which allows the user to generate and compare as many predictor maps as they want without re-fitting the model.

Simulations further demonstrated the interpretability advantage of T-PLS predictors when the “ground truth” is known. We simulated a 2D brain signal on 17×17 voxel grid, in which only the center 5×5 grid had signal predictive of Y (**Fig. 7A**). This image was subsequently smoothed to simulate smoothness of fMRI images (**Fig. 7B**). When we fit a OLS predictor on this simulated data, it highlighted the canonical problem with fMRI images: multicollinearity. OLS predictors resulted in voxel coefficients that were tessellating in alternating signs, which makes it impossible to know whether the signal is positively predictive or negatively predictive (**Fig. 7C**). LASSO deals with this multicollinearity by selecting only those variables that are the most useful in prediction and removing the rest. The resulting predictor, therefore, is very sparse and it is difficult to identify the regional pattern (**Fig. 7D**). PCR-LASSO uses PCA data-reduction to create locally smooth predictors that reflect the smoothness of fMRI images. However, there is no voxel selection, making it more difficult to distinguish regions that are predictive from those that are not, and the predictor contains PCA-based artifacts (**Fig. 7E**; negative coefficients on the edges of the image). Of all these methods, T-PLS provides a predictor that most closely resembles the ground-truth signal, detecting the positively predictive 5×5 signal grid in the center and eliminating all other voxels from the predictor. These clustered coefficients help the researcher the regions that are predictive from those that are not.

**Figure 7.**
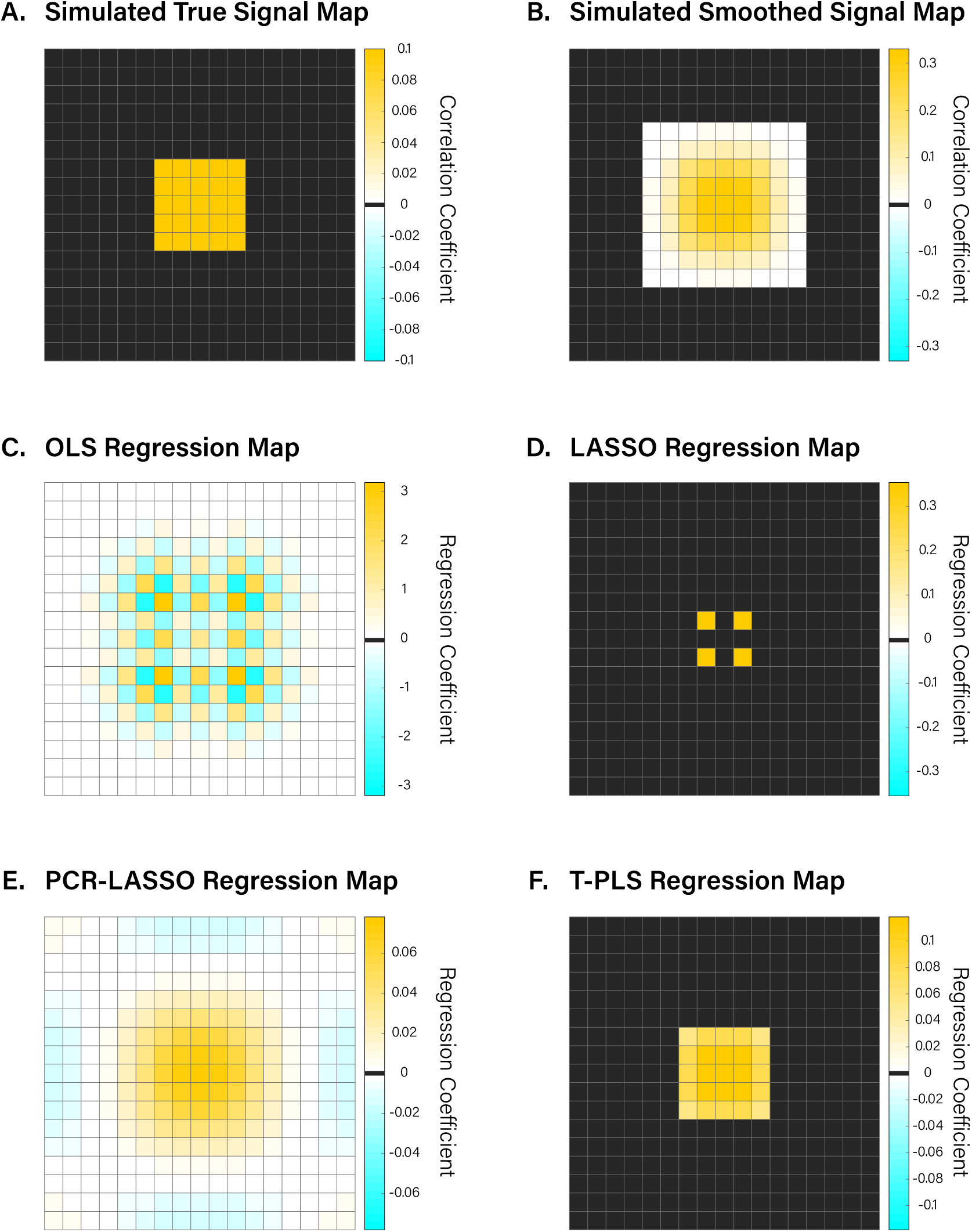
Simulated neuroimaging signal and various predictor fits. Panels A shows a simulated 17×17 voxel grid where only the center 5×5 grid has signal that is predictive of Y at correlation *r* = 0.1. Panel B shows the result of a spatial smoothing filter to panel A, meant to simulate fMRI image smoothness. Coefficients that are exactly 0 were marked as black, while those that are close to 0 are marked as near white. Panel C~F shows regression coefficients from OLS regression, LASSO regression, PCR-LASSO regression, and T-PLS, respectively. Only panels D and F have variable selection, thereby making most voxels exactly 0 (marked black).

## Discussion

There is growing interest in using neuroimaging to build predictors that can decode mental states or behavior. Whole brain decoding methods promise several advantages over partial-brain methods as they can harness more signal across the brain, identify unique neural signatures, and illuminate relationships between brain regions. However, methods for whole-brain prediction face several challenges in terms of computation time, memory usage, and, most importantly, interpretability. To this end, we present here a novel method, thresholded partial least squares (T-PLS), that provides computationally feasible, interpretable, whole-brain predictors. T-PLS exploits analytical properties of partial least squares (PLS) algorithms to dramatically reduce model fitting time, use less computational memory, and still provide high predictive performance. Compared in a real neuroimaging dataset against extant methods PCR-LASSO and LASSO, T-PLS was up to 25 times faster and used as much as 80% less computation memory. Furthermore, T-PLS was the only method whose per-subject fitting time was scalable and did not increase as the dataset size increased. T-PLS also showed higher out-of-sample predictive performances than other whole-brain methods.

T-PLS builds upon previous uses of partial least squares (PLS) in fMRI prediction (Kragel & LaBar, 2014; McIntosh et al., 1996) by introducing variable (voxel) selection that is based on fast analytical computation and cross-validation. By calculating the PLS components’ z-statistics, which are usually not needed since PLS components are created to explain the most covariance, T-PLS approximates the z-statistics of each voxel by back-projection and hence analytically computes their relative importance. This is an improvement from previous thresholding approaches which either used arbitrary threshold for the sake of interpretability (McIntosh et al., 1996), or used time-consuming bootstrap measures to calculate each voxels’ p-values and create a thresholded map (Kragel & LaBar, 2014), which are again different from the actual map that was used for prediction.

Furthermore, unlike PCR-LASSO, where LASSO was used to select the most useful PCA components, T-PLS’s component selection method yields the same result as LASSO selection without having to fit LASSO. This is because in LASSO regression, if all predictor variables are orthogonal, the variable selection order follows the absolute size of the coefficients, which in the case of PLS, coincides with the original order of PLS components, since components are extracted in the order of most covariance explained.

These two features of variable selection and component selection allow T-PLS to boast a unique ‘fit-once-tune-later’ feature, where the model is fit only once and the two tuning parameters are chosen after the fact. This is in stark contrast to other modern prediction algorithms where models need to be re-fit for every tuning parameter combination. Not only does this allow for faster cross-validation, but it also allows researchers to explore various tuning parameters to determine which brain regions are important for prediction and can choose the best level of sparsity given the tradeoff between parsimony and performance.

We acknowledge that principal component regression (PCR) and LASSO, since they are widely used methods outside of neuroimaging, have a large number of potential addendums and improvements available. For example, there are stochastic variants of PCA that can deal with the huge RAM and computation time costs of PCA for large datasets (Halko, Martinsson, & Tropp, 2011). These variants may reduce the computation time and RAM usage of the PCR-LASSO method shown in this paper, but it still stands that the principal components may not be the best suited for prediction and that the resulting whole-brain predictor is less interpretable than T-PLS. For LASSO, there are stochastic gradient descent-based methods that can significantly reduce the model fitting time by evaluating the likelihood function on subsamples of the data. These methods, however, face a tradeoff with the accuracy of the fit as the gradient calculations are approximations. Concordantly, we found that these methods were both slower and less accurate than T-PLS, and we have excluded them from this paper.

It is also important to acknowledge that we did not compare all existing whole-brain prediction methods. For example GraphNet was proposed as a purely likelihood-based method that can yield interpretable whole-brain predictors using modified elastic-net regression (Grosenick et al., 2013). While GraphNet is a principled method of dealing with whole-brain prediction, given that the penalties are based on elastic-net, which has more tuning parameters than LASSO, we expect this algorithm to be slower than the LASSO algorithm that we compared in this manuscript. Furthermore, previous research has found that while GraphNet yields interpretable predictors, it does so at the cost of predictive performance compared to LASSO (Mohr, Wolfensteller, Frimmel, & Ruge, 2015), which in this manuscript showed lower performance than T-PLS. We also did not compare support vector machines as we expect it to be similar to LASSO in result (if L1-penalized). Finally, we also left out non-linear methods such as neural nets (Thomas, Heekeren, Müller, & Samek, 2019), gaussian processes (Marquand et al., 2010), or naïve-bayes (Kassam et al., 2013) as they are expected to be considerably slower, with many more tuning parameters, and may require a different method of interpreting the resulting predictors.

We see the relationship between whole-brain prediction methods and ROI-based prediction methods as similar to that between multivariate regression and pairwise bivariate correlation. ROI-based methods may yield insight about how one specific region is related to a mental process, but a whole-brain method can yield insight about how multiple ROIs relate to one another and contribute to prediction; they are both important analysis tools that every researcher must use to understand the whole picture. Concordantly, we see whole-brain prediction as an important analysis tool in the coming years of neuroimaging, and we hope that the method that we propose here, along with the provided packages, can make this analysis a convenient and essential part of neuroimaging analysis pipelines.

## Supplemental Materials

### Neuroimaging data quality control

Some of the data from Kable et al. (2017) was removed from the analysis due to trivial reasons. To keep the number of observations per subject roughly similar, we excluded four pilot participants who had more trials than others. Counting both session 1 and session 2 data, we had 286 sessions worth of data, each with 120 binary choices. From here, we removed 4 intertemporal choice sessions and 6 risky choice sessions that had premature termination of scan due to technical issues. Additionally, 13 intertemporal choice sessions were removed for having extremely unbalanced choices (either accept or reject more than 95% of the time), and 6 sessions were removed for having too many missed responses (more than a quarter worth of session). For risky choice, 10 sessions were removed for unbalanced choices and 6 sessions were removed for too many missed responses. In total, we had 264 sessions worth of data for ITC and 267 sessions worth of data for RC. While most participants had both ITC and RC tasks, since several subjects only had one task, we decided to treat these two tasks’ sessions as separate participants for our analyses.

### Image preprocessing and single trial deconvolution

The Kable et al., (2017) dataset was acquired with a Siemens 3T Trio scanner with a 32-channel head coil. High-resolution T1-weighted anatomical images were acquired using an MPRAGE sequence (T1 = 1100ms; 160 axial slices, 0.9375 × 0.9375 × 1.000 mm; 192 × 256 matrix). T2*-weighted functional images were acquired using an EPI sequence with 3mm isotropic voxels, 64 × 64 matrix, TR = 3,000ms, TE = 25ms, 53 axial slices, 104 volumes. B0 fieldmap images were collected for distortion correction (TR = 1270ms, TE = 5 and 7.46ms). The images were preprocessed via fMRIPrep. The preprocessing pipeline, in short, performed motion-correction, slice-time correction, and b0-map unwarping on all runs and registered and resampled to a MNI 2mm template. The authors of fMRIPrep has requested the automatically generated preprocessing info to be pasted into the manuscript in its unaltered form. Given its length, we provide them in the supplemental materials.

For estimating the activity of each trial, we used beta-series regression (Rissman, Gazzaley, & D’Esposito, 2004). The regressors were time-locked to the trial onset period with event duration of 0.1 seconds and convolved with a gamma HRF function. The last trial of each run was excluded from analysis because the BOLD activity of the last trial was often not observed due to the termination of the scan. This gave us 29 regressor of interest per 1 run of scan. Additionally, we included the following nuisance regressors which were generated from fmriprep: cosine components for high-pass filtering, CSF signal, white matter signal, global signal, standard 6 motion regressors, and 6 PCA components from an anatomical mask of white matter and CSF (‘a_comp_cor’). After the single trial coefficients were estimated, all images were smoothed with a FWHM 5mm gaussian filter. To make analysis easy, we only used the voxels that were active for all subjects; this gave us a fairly conservative mask of the brain with 184,319 voxels.

### Fmriprep boilerplate

Results included in this manuscript come from preprocessing performed using fMRIPrep 20.0.5 (Esteban, Markiewicz, et al. (2018); Esteban, Blair, et al. (2018); RRID:SCR_016216), which is based on Nipype 1.4.2 (Gorgolewski et al. (2011); Gorgolewski et al. (2018); RRID:SCR_002502).

#### Anatomical data preprocessing

A total of 2 T1-weighted (T1w) images were found within the input BIDS dataset. All of them were corrected for intensity non-uniformity (INU) with N4BiasFieldCorrection (Tustison et al. 2010), distributed with ANTs 2.2.0 (Avants et al. 2008, RRID:SCR_004757). The T1w-reference was then skull-stripped with a Nipype implementation of the antsBrainExtraction.sh workflow (from ANTs), using OASIS30ANTs as target template. Brain tissue segmentation of cerebrospinal fluid (CSF), white-matter (WM) and gray-matter (GM) was performed on the brain-extracted T1w using fast (FSL 5.0.9, RRID:SCR_002823, Zhang, Brady, and Smith 2001). A T1w-reference map was computed after registration of 2 T1w images (after INU-correction) using mri_robust_template (FreeSurfer 6.0.1, Reuter, Rosas, and Fischl 2010). Volume-based spatial normalization to one standard space (MNI152NLin2009cAsym) was performed through nonlinear registration with antsRegistration (ANTs 2.2.0), using brain-extracted versions of both T1w reference and the T1w template. The following template was selected for spatial normalization: ICBM 152 Nonlinear Asymmetrical template version 2009c [Fonov et al. (2009), RRID:SCR_008796; TemplateFlow ID: MNI152NLin2009cAsym],

### Functional data preprocessing

For each of the 18 BOLD runs found per subject (across all tasks and sessions), the following preprocessing was performed. First, a reference volume and its skull-stripped version were generated using a custom methodology of fMRIPrep. A B0-nonuniformity map (or fieldmap) was estimated based on a phase-difference map calculated with a dual-echo GRE (gradient-recall echo) sequence, processed with a custom workflow of SDCFlows inspired by the epidewarp.fsl script and further improvements in HCP Pipelines (Glasser et al. 2013). The fieldmap was then co-registered to the target EPI (echo-planar imaging) reference run and converted to a displacements field map (amenable to registration tools such as ANTs) with FSL’s fugue and other SDCflows tools. Based on the estimated susceptibility distortion, a corrected EPI (echo-planar imaging) reference was calculated for a more accurate co-registration with the anatomical reference. The BOLD reference was then co-registered to the T1w reference using flirt (FSL 5.0.9, Jenkinson and Smith 2001) with the boundary-based registration (Greve and Fischl 2009) cost-function. Co-registration was configured with nine degrees of freedom to account for distortions remaining in the BOLD reference. Head-motion parameters with respect to the BOLD reference (transformation matrices, and six corresponding rotation and translation parameters) are estimated before any spatiotemporal filtering using mcflirt (FSL 5.0.9, Jenkinson et al. 2002). BOLD runs were slice-time corrected using 3dTshift from AFNI 20160207 (Cox and Hyde 1997, RRID:SCR_005927). The BOLD time-series (including slice-timing correction when applied) were resampled onto their original, native space by applying a single, composite transform to correct for head-motion and susceptibility distortions. These resampled BOLD time-series will be referred to as preprocessed BOLD in original space, or just preprocessed BOLD. The BOLD time-series were resampled into standard space, generating a preprocessed BOLD run in MNI152NLin2009cAsym space. First, a reference volume and its skull-stripped version were generated using a custom methodology of fMRIPrep. Several confounding time-series were calculated based on the preprocessed BOLD: framewise displacement (FD), DVARS and three region-wise global signals. FD and DVARS are calculated for each functional run, both using their implementations in Nipype (following the definitions by Power et al. 2014). The three global signals are extracted within the CSF, the WM, and the whole-brain masks. Additionally, a set of physiological regressors were extracted to allow for component-based noise correction (CompCor, Behzadi et al. 2007). Principal components are estimated after high-pass filtering the preprocessed BOLD time-series (using a discrete cosine filter with 128s cut-off) for the two CompCor variants: temporal (tCompCor) and anatomical (aCompCor). tCompCor components are then calculated from the top 5% variable voxels within a mask covering the subcortical regions. This subcortical mask is obtained by heavily eroding the brain mask, which ensures it does not include cortical GM regions. For aCompCor, components are calculated within the intersection of the aforementioned mask and the union of CSF and WM masks calculated in T1w space, after their projection to the native space of each functional run (using the inverse BOLD-to-T1w transformation). Components are also calculated separately within the WM and CSF masks. For each CompCor decomposition, the k components with the largest singular values are retained, such that the retained components’ time series are sufficient to explain 50 percent of variance across the nuisance mask (CSF, WM, combined, or temporal). The remaining components are dropped from consideration. The head-motion estimates calculated in the correction step were also placed within the corresponding confounds file. The confound time series derived from head motion estimates and global signals were expanded with the inclusion of temporal derivatives and quadratic terms for each (Satterthwaite et al. 2013). Frames that exceeded a threshold of 0.5 mm FD or 1.5 standardised DVARS were annotated as motion outliers. All resamplings can be performed with a single interpolation step by composing all the pertinent transformations (i.e. head-motion transform matrices, susceptibility distortion correction when available, and co-registrations to anatomical and output spaces). Gridded (volumetric) resamplings were performed using antsApplyTransforms (ANTs), configured with Lanczos interpolation to minimize the smoothing effects of other kernels (Lanczos 1964). Non-gridded (surface) resamplings were performed using mri_vol2surf (FreeSurfer).

## Footnotes

1 Technically, the calculation of the components and their regression coefficients are done simultaneously based on analytical methods without having to perform regression

2 The original GLMNET package for MATLAB could not import a dataset size of as large a magnitude as in this study because the FORTRAN API with MATLAB was written in 32-bit architecture; we have updated the FORTRAN code ourselves to 64-bit architecture to circumvent this issue; the updated package is provided here: https://github.com/sangillee/GLMNET64MATLAB

